# McSNAC: A software to approximate first-order signaling networks from mass cytometry data

**DOI:** 10.1101/2021.12.02.470955

**Authors:** Darren Wethington, Sayak Mukherjee, Jayajit Das

## Abstract

Mass cytometry (CyTOF) gives unprecedented opportunity to simultaneously measure up to 40 proteins in single cells, with a theoretical potential to reach 100 proteins. This high-dimensional single-cell information can be very useful to dissecting mechanisms of cellular activity. In particular, measuring abundances of signaling proteins like phospho-proteins can provide detailed information on the dynamics of single-cell signaling processes. With a proper computational analysis, timestamped CyTOF data of signaling proteins could help develop predictive and mechanistic models for signaling kinetics. These models would be useful for predicting the effects of perturbations in cells, or comparing signaling networks across cell groups. We propose our Mass cytometry Signaling Network Analysis Code, or **McSNAC**, a new software capable of reconstructing signaling networks and estimating their kinetic parameters from CyTOF data.

McSNAC approximates signaling networks as a network of first-order reactions between proteins. This assumption breaks down often as signaling reactions can involve binding and unbinding, enzymatic reactions, and other nonlinear constructions. Furthermore, McSNAC may be limited to approximating indirect interactions between protein species, as cytometry experiments are only able to assay a small fraction of the protein species that are involved in signaling. We carry out a series of *in silico* experiments here to show that 1) McSNAC is capable of accurately estimating the ground-truth model in a scalable manner when given data originating from a first-order system; 2) McSNAC is capable of qualitatively predicting outcomes to perturbations of species abundances in simple second-order reaction models and in a complex *in silico* nonlinear signaling network in which some proteins are unmeasured. These findings demonstrate that McSNAC can be a valuable screening tool for generating models of signaling networks from timestamped CyTOF data.

## Introduction

Single cells respond to stimulus via receptors that are usually bound to the plasma membrane. These receptors bind to cognate ligands generated by a stimulation, such as viral infection of cells in the local environment, and initiate a series of biochemical signaling reactions leading to activation of many genes that can generate a variety of cell responses such as secretion of specific proteins (e.g., cytokines), cell proliferation, or cell death [1-3]. Signaling reactions are composed of biochemical reactions involving a large number of proteins (∼thousands), and it is often challenging to determine key signaling regulators of specific cell responses among a large set of interacting proteins [4, 5]. This problem is further confounded as single cells can contain large cell-cell variations of protein abundances where protein abundances are usually distributed broadly (e.g., lognormally) in a cell population, even in a clonal population[6]. Since biochemical reaction propensities depend on concentrations of participating molecules, and biochemical reactions are intrinsically stochastic in nature due to thermal fluctuations[7], single cells in clonal cell populations stimulated by the same ligand can show large variations in responses making it difficult to determine key regulators in the face of large variations. Recent advances in single cell cytometry, such as Mass Cytometry, or Cytometry by Time-of-Flight (CyTOF), are currently capable of measuring up to 40 proteins simultaneously in thousands to millions of single cells, with a theoretical limit as high as 100 proteins[8, 9]. Thus, measuring signaling protein abundances at different points post stimulation using CyTOF provides time stamped snapshot data regarding signaling kinetics and cell-cell variation of the kinetics. However, it is difficult to intuitively determine key regulators that give rise to a specific response in a single cell or a group of single cells with this data alone for the following reasons. First, though causal relationships between measured proteins, such as phosphorylation of protein A inducing phosphorylation of protein B, might be known from previous experiments, activation of key regulator proteins can often be produced by multiple proteins interacting synergistically. Second, protein abundances vary widely across cells, and proteins in the same individual cell are not measured across time, thus if specific proteins are activated during signaling kinetics in individual cells remains unknown.

Some of the above challenges are also present in deciphering mechanisms underlying signaling kinetics using population data or traditional flow cytometry data measuring few proteins. Mechanistic mathematical models composed of ordinary differential equations (ODEs) or stochastic models describing biochemical signaling reactions have been used to address these difficulties[10, 11]. These models work well when the number of protein species is small and biochemical reaction propensities for reacting proteins are well characterized. However, it is often not the case for newly discovered cell types (e.g., MDSCs)[12] or situations where multiple types of receptors (e.g., cytokine receptors and primary cell receptor) synergize[13]. It is challenging to build such mechanistic models in such situations where protein species interacting physically are usually not measured simultaneously and it is unclear what form (e.g., first, second or higher order reaction) of reaction propensities should be used for describing the interaction between two protein species separated by many intermediate reactions.

Machine learning has also been employed to characterize ODEs to predict the effects of cellular perturbations[14]. This model is quite useful for mechanistically predicting the effects of perturbations, but does not have a mechanism to incorporate the single-cell resolution offered by CyTOF. It is important to note that signaling processes happen at the single-cell resolution, and thus single-cell experiments contain crucial information on signaling dynamics. This information can be used to identify how cells vary in their protein expressions, and which proteins co-vary with each other in single cells. For these reasons, it is crucial to have a method to form accurate mechanistic models for single-cell signaling networks.

To this end, we developed a first order mass action reaction kinetics model given by linear ordinary differential equations (ODEs) to describe signaling kinetics over a time interval – e.g., time between two successive mass cytometry measurements[13, 15]. The solution of the model describes the time evolution in a closed form and accounts for cell-cell variations of protein abundances. In this paper we report development of a software package Mass cytometry Signaling Network Analysis Code (McSNAC) that takes flow cytometry data at two successive time points as input, fits the model involving first order kinetics, and generates parameter estimates for the kinetic rates. The model can then be used for generating predictions regarding changes in protein abundances. We use McSNAC to model synthetic cytometry data to demonstrate: 1. Parameters in the model can be well estimated. 2. Models that include the ground truth model describing first order reaction kinetics provide better fits than models without the ground truth. 3. The first order models in McSNAC are able to make accurate qualitative predictions to perturbations in protein abundances. 4. The models in McSNAC generate better predictions to perturbations when directionality of the underlying reaction is captured in the model.

## Model

McSNAC describes kinetics of single cell protein abundances using first order chemical reaction kinetics. Consider *p* number of protein species {X_1_,⋅ ⋅ ⋅, X_*p*_} measured in cytometry experiments that interact during a signaling process. In McSNAC the abundance *x*^{*α*}^_*i*_ of protein X_i_ in each cell indexed by α changes with time given by the linear ODEs shown below:

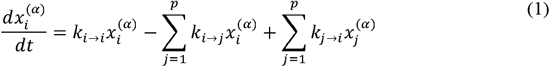

where the first order rate that produces species X_j_ from X_i_ is given by *k*_*i* → *j*_ (*k*_*i*→ *j*_ ≥0 for i≠j). *k*_*i*→ *i*_ >0 denotes self-production or production of X_i_ from another unmeasured species while *k*_*i* →*i*_<0 denotes self-decay. In the above kinetics, the kinetic rates {*k*_*i* → *j*_} are the same across single cells (i.e., a *k*_*i* → *j*_ does not depend on the cell index α). This is an approximation as some of these rates could represent nonlinear reaction such as enzymatic modifications where the rates could depend on the abundances of an enzyme which could vary from cell-to-cell. The ODEs in Eq. 1 can be represented in compact form by introducing a *p×p* matrix, *M*, where *M*_*ij*_ (*i* ≠ *j*) = *k*_*j* → *i*_, and *M*_*ii*_ = *k*_*i* → *j*_ − Σ_*j*_ *k*_*i* → *j*_ =, i.e.,

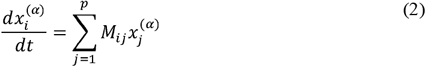

This representation of the mass-action kinetics offers a convenient closed-form solution for 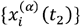 at given 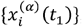, i.e.,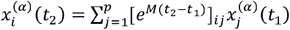.

Mass cytometry experiments are unable to track single cells over time as individual cells are destroyed at the time of measurement. However, it is still possible to follow the kinetics of average values, covariances and higher order moments of the protein abundances that are computed from the snapshot data.

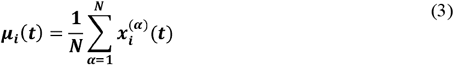

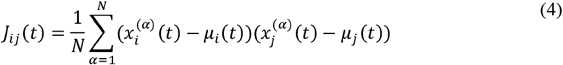

The ODEs in Eq. (2) can be solved to relate the average values {μ_i_} and covariances {J_ij_} calculated at times t_1_ and t_2_,

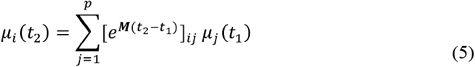

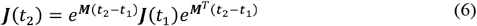

The bold form symbols in the above equations denote matrices. These above closed-form solutions make it efficient to compute predicted average values and covariances in the model compared to iteratively solving the ODEs in Eq. (2) numerically and then computing those variables using Eqs. (3)-(4). Given the measured values of {μ_i_}and {J_ij_} at time t_1_ from cytometry experiments we estimated the kinetic rates in *M* that best fit cytometry data for {μ_i_}and {J_ij_} at a later time t_2_. *M* is estimated by minimizing a cost function (Eq. 7) by employing a simulated annealing algorithm. The cost function is defined as,

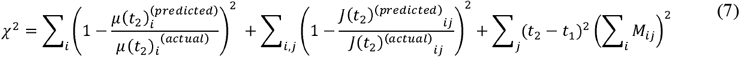

Where, μ(t_2_)^(actual)^ and J(t_2_)^(actual)^ are the means and covariances calculated from cytometry data at time t_2_ using Eqs. (3) and (4), and, μ(t_2_)^(predicted)^ and J(t_2_)^(predicted)^ are the means and covariances predicted by the model following Eqs. (5) and (6) given μ(t_1_) and J(t_1_) calculated from cytometry data. This cost function has three terms, penalizing different aspects of the fit. The first term penalizes deviations from the observed averages. The second term penalizes deviations from the observed variances and covariances. The third term penalizes kinetics that do not abide by conservation of mass (assuming phospho-groups are transferred and no other changes in abundance occur). In other words, the third term penalizes *k*_*i* → *i*_ ≠ 0 from Eq. 1. If data come from a ground-truth first-order model which abides by conservation of mass, and the ground-truth *M* is found, then χ^2^ =0. A more detailed description of this method can be found in Mukherjee et al. 2017[13].

## Results

### McSNAC is capable of accurately estimating first-order reaction rates

Here we describe McSNAC’s ability to (i) model data when the wiring of the ground truth model is known but the values of the rate constants are unknown, (ii) computationally scale as the number of proteins in the data is increased up to 40, and, (iii) separate models that do or do not include the ground truth. The synthetic cytometry data were generated for a coupled set of first-order reactions as shown in Reaction R1 using Eq. 2, where the distributions of {x_1_^(α)^, .., x_p_^(α)^} at t=0 are chosen from log-normal distributions (see Materials and Methods for details).

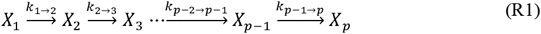

The rates {*k*_*i* → *j*_} were assigned from a uniform random distribution with a range of 0 to 1, and {x_1_^(α)^, .., x_p_^(α)^} were computed for 2500 single cells at two different time points. This constituted the synthetic cytometry data which were generated for a range of *p* between 8 to 40. We applied McSNAC to synthetic data following the architecture in R1 to determine (i) how accurately parameters {*k*_*i* → *j*_} are estimated, (ii) how does the run time increase with increasing *p*. The comparison of the McSNAC estimated parameters and their actual values shows high accuracy in parameter estimation (Fig 1a, 1b). The run time of McSNAC on a single CPU (Intel 2.0 GHz, Xeon E5-2640v2, compiled with Intel Fortran compiler *ifort*) ranged from minutes to hours where the case for *p*=40 took 11.9 hours to complete on average (Fig 1c). The dependence of the run time with *p* varies as ∼ *p*^3^ indicating that this technique is scalable with the large number of inputs afforded by CyTOF. It is important to note that exact runtimes vary depending on hyperparameter specification, such as number of temperature cooling iterations and number of Monte Carlo samples per temperature step, and the exact data and model being fitted.

**Fig 1:**
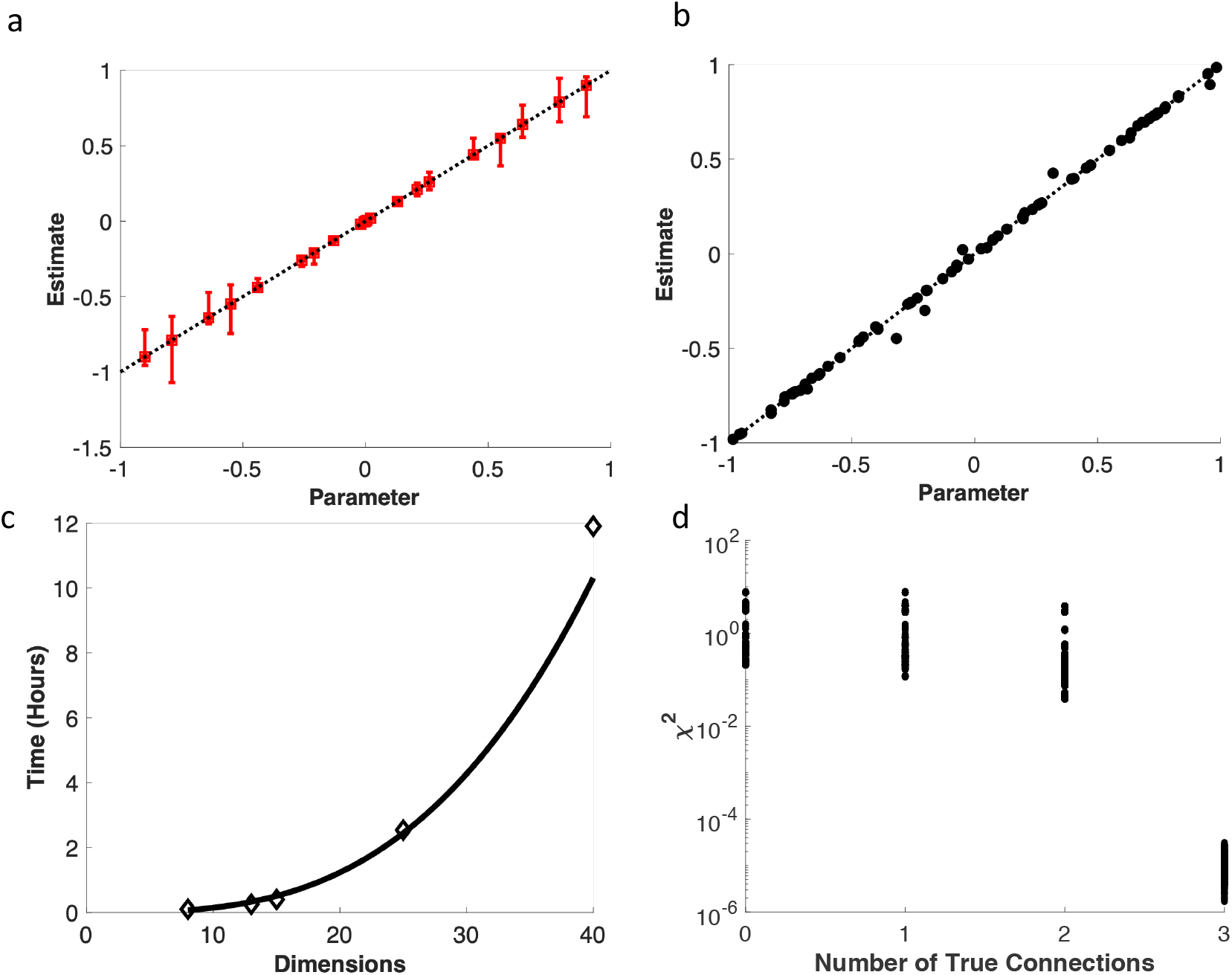
McSNAC is a scalable software capable of reconstructing first-order signaling dynamics. a) Comparison of manually selected parameters of a 13-dimensional system and the estimated parameter values from McSNAC. Error bars denote confidence intervals on the parameter estimates. The dotted black line represents y=x, and perfectly estimated parameters will lie on this line. b) Comparison of manually selected parameters of randomly selected parameters of a 40-dimensional system and the estimated value of each parameter by McSNAC. The dotted black line represents y=x, and perfectly estimated parameters will lie on this line. c) The average time (n=3) required to run McSNAC for the number of dimensions *p* shown for a system represented by Equation R1. d) 2^12^ models were simulated, each one containing either none (0), some (1-2), or all (3) of the k_i→j_ connections in the ground truth model. Models with all (3) of the true connections, comprising the ground truth model, generate χ^2^≅0, indicating a perfect fit.

Next, we evaluated the ability of the cost function (Eq. 7) to separate models that include or do not include the architecture of the ground truth model. This is relevant for cases where interactions between protein species in a signaling network are not known and several potential network architectures can be hypothesized to describe the data. To test the above property, we generated synthetic cytometry data for a reaction scheme involving four protein species given by *p*=4 in R1. If we had no knowledge of the architecture of the ground truth network that connected the proteins, these four proteins would offer a total of 12 possible *k*_*i* → *j*_’s, and 2^12^ unique network architectures (more details in Materials and Methods section). Applying McSNAC to each of the 2^12^ unique network architectures and determining the goodness-of-fit χ^2^ shows that when a model contained the ground truth architecture it generated close to zero values of χ^2^, whereas models that do not include the ground truth architecture generated larger values of χ^2^ (Fig 1d). These results show that McSNAC is capable of reconstructing the ground truth signaling dynamics from data resulting from a first-order system, regardless of dimensionality or *a priori* information of its wiring.

In the next sections we show results when McSNAC is applied for data sets that represent scenarios that are commonly present in cytometry datasets studying signaling kinetics. Signaling biochemical reactions contain reactions associated with propensities that are nonlinear functions of abundances of reacting protein species. Moreover, a large number (∼ 1000-10000) of proteins and their chemically modified forms can be involved in signaling reactions, and CyTOF experiments are able to measure a small fraction of those protein species. Therefore, we tested the ability of the first order reaction kinetics in McSNAC to approximate the signaling reaction kinetics in the above situations. The following cases are considered: (i) The ground truth model contains non-linear biochemical reactions. (ii) Not all proteins participating in biochemical signaling reactions are measured. We find that McSNAC is able to describe the result of perturbations such as increase and decrease of specific protein abundances in most situations, however, McSNAC performs poorly when it comes to predicting protein abundances at a later time (Fig. S1). In the next sections we provide further details regarding McSNACs ability to predict outcomes of perturbations of signaling kinetics due to increase and decrease of specific protein abundances. Such perturbations are routinely performed using specific drugs such as Src family kinase inhibitor, Syk family kinase inhibitors, or small interfering RNA (siRNA).

### McSNAC is capable of approximating simple non-linear systems

We test the ability of McSNAC to capture kinetics in a ground truth model describing a second-order reaction in which two reactants combine to form a product. The candidate models probed by McSNAC are composed of first-order reactions. We generated *in silico* data formed by ground truth model GT1 given by Reaction R2 at two different time points and fitted the candidate models (C1 to C7) shown in Table 1 to estimate reaction rates.

**Table 1:**
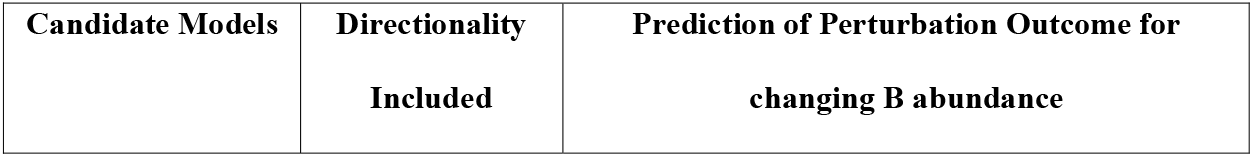
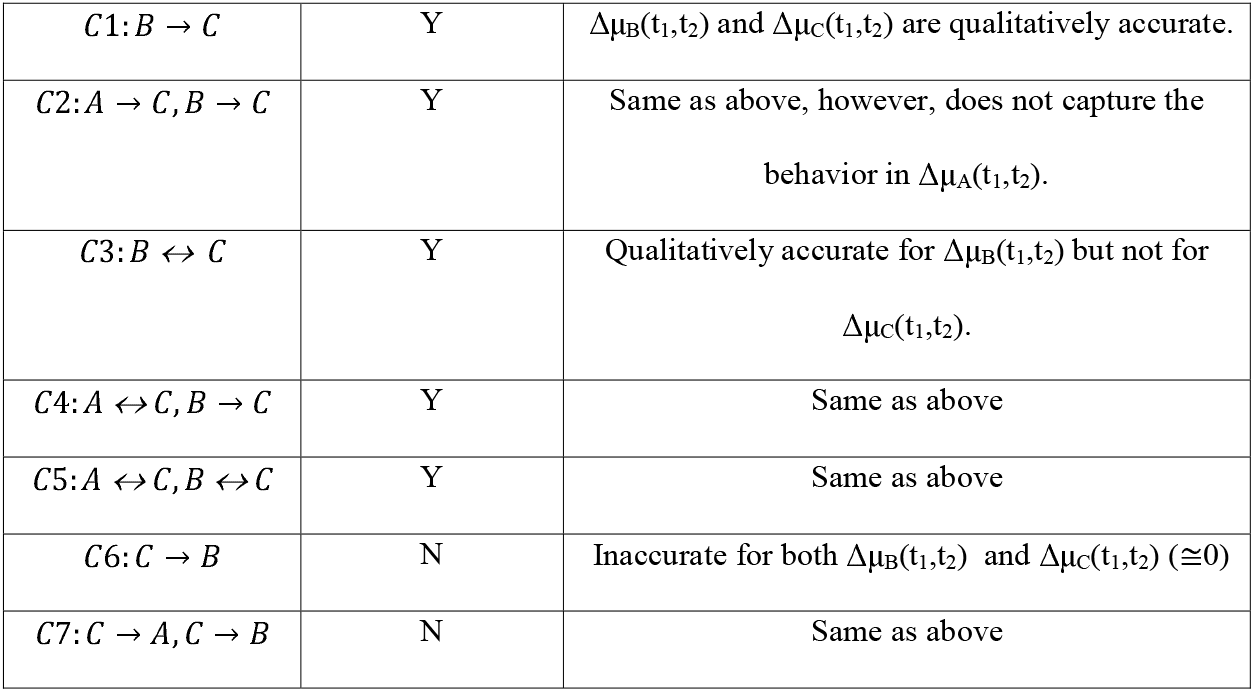
List of candidate first order models for describing a unidirectional second-order reaction A+B → C in ground truth model GT1.

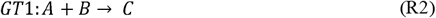

The McSNAC candidate models with the best fit *M* values describe few abundances at t_2_ reasonably well, but over- or undershot-values for certain species. For example, candidate model C1 (B→ C) matches mean abundances of B but overestimates the mean abundance of C for model GT1 at t_2_ (details in the Supplementary Material). This behavior is expected as McSNAC uses first order reactions to approximate second order reactions in the ground truth model. Next, we evaluated how the candidate models in McSNAC perform for predicting outcomes of in silico perturbation experiments where increased abundances of species (e.g., B) at t_1_ and assessed the ability of the candidate models to predict changes in the abundances of all the species at a later time t_2_. We defined a variable ΔμX(t_1_,t_2_) describing changes in mean abundances of species X between two successive times t_1_ and t_2_(>t_1_) for quantifying comparisons between the ground truth and candidate models in the perturbation experiments. ΔμX(t_1_,t_2_) is given by,

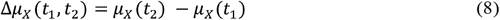

ΔμX (t_1_,t_2_) > 0 (or <0) indicates net production (or consumption) in the mean abundance of X as the kinetics progress from time t_1_ to t_2_. To illustrate how ΔμX (t_1_,t_2_) can be used to analyze outcomes of a perturbation experiment, if ΔμC (t_1_,t_2_) > 0 in the ground truth model and also in a candidate model (e.g., C1) when abundance of B is increased at time t_1_, it implies that candidate model C1 is able to qualitatively predict the outcome of the perturbation. The quantitative difference between the ground truth model and the prediction made by the candidate model can be given by the squared distance between ΔμX (t_1_,t_2_) obtained from the ground truth and a candidate model. First, we investigated results from increasing or decreasing abundances of B at time t_1_ in the ground truth model GT1 and the candidate models (Table 1, Fig. 2a, Fig. S2). The candidate models C1 and C2 correctly predict the net production of species B and C or ΔμB (t_1_,t_2_) > 0 and ΔμC (t_1_,t_2_) > 0 as in GT1. In addition, as abundance of B is increased (or decreased) in the perturbation experiment the productions of B and C are increased (or decreased); this monotonic behavior is also correctly predicted by candidate models C1 and C2. However, the values of ΔμB (t_1_,t_2_) and ΔμC (t_1_,t_2_) in the models C1 and C2 are different than that in the ground truth model GT1 indicating that the candidate models are able to correctly predict outcomes of perturbations qualitatively but not quantitatively (Table 1 and Fig. 2a). The models C3 and C7 are unable to correctly predict the above qualitative changes correctly (Table 1 and Fig. 2a). However, none of the candidate models are able to capture the decrease (or increase) in ΔμA(t-_1_,t_2_) as abundance of B is increased (or decreased) in GT1 (Fig. 2b) as the first order reactions are unable to capture the dependencies of consumption of A and B given by the second order reaction in GT1. Thus, as long as the directionality of the reaction, i.e., irreversible production of C from B and A is correctly captured in the candidate models they are able to qualitatively predict outcomes of the perturbation for most of the species.

**Fig 2:**
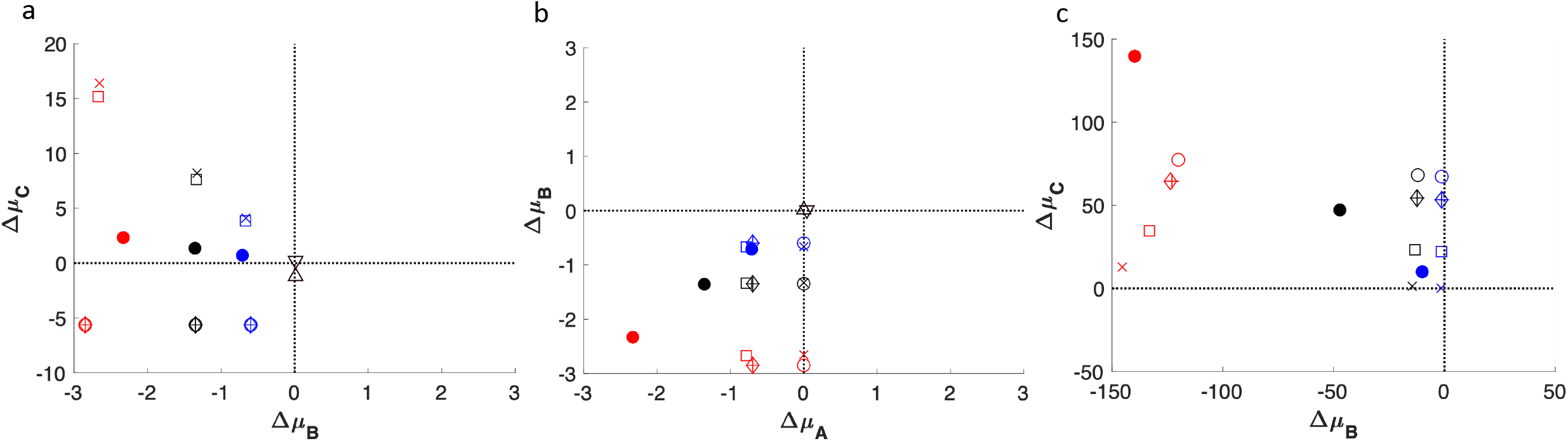
McSNAC is capable of predicting the effects of perturbations in simple nonlinear systems. a) Predictions of the change in averages through time in B and C resulting from perturbations of B for the ground-truth model indicated by Equation GT1, as predicted by the approximations in Equations C1-C7. Large, filled circles represent the ground truth, and markers for candidate predictions are as follows. C1: *B* → *C* (X’s); C2: *A* → *C, B* → *C* (Squares); C3: *B* ↔ *C* (Open Circles); C4: *A* ↔ *C,B* → *C* (Diamonds); C5: *A* ↔ *C,B* ↔ *C* (+’s); C6: *C* → *B* (upward-facing triangles); and C7: *C* → *A,C* → *B* (downward-facing triangles). Black points are estimations without a perturbation, red points are estimations in which B was perturbed upwards two-fold, and blue points are estimations in which B was perturbed downwards two-fold. Δμ_B_ and Δμ_C_ are defined as the change in averages through time, shown in Equation 8. b) Predictions of the change in averages through time in A and B resulting from perturbations of B for the ground-truth model indicated by Equation GT1. Markers and perturbation information are the same as in (a). c) Predictions of the change in averages through time resulting from perturbations of B for the ground-truth model indicated by Equation GT2, as predicted by the approximations in Equations C1-C5. Markers are the same as in (a), but perturbations of B are 10-fold.

To extend the exploration of approximating simple nonlinear dynamics, we simulated data from a similar nonlinear system shown by Reaction R3.

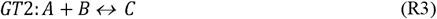

Because the directionality of this model depends on the relative abundance of reactants and products, we formed three sets of data: 1) Reactants are more abundant than product (rightward directionality), 2) Product is more abundant than reactants (leftward directionality), 3) Reactants and product are approximately equal in magnitude (almost no directionality). We modeled each set of data formed by this nonlinear system with the approximations in candidate models C1-C5. We show that for the first case (rightward directionality) we can qualitatively capture the predicted effects of a perturbation of B (Fig. 2c). However, perturbations of A predicted larger effects in ΔμC for these data than for GT1 (Fig S3). An important note is that perturbations of C again generated no impact on either ΔμA or ΔμB, which is not consistent with the bidirectional nature of GT2. This could be explained by the net forward directionality of the data construction, where a net production of C occurs when no perturbation occurs. A more complete description of the results of models C1-C5 on each dataset is provided in Supplementary Table 1.

### McSNAC is capable of predicting effects of perturbations in complex non-linear signaling networks

Here we applied McSNAC to test its ability to predict effects of changes in species abundances for a signaling model describing membrane proximal signaling events in an NK cell. The signaling reaction contains multiple binding-unbinding and enzymatic reactions – all of which are composed of coupled second order reactions. The NK cell signaling model, which is the ground truth model here, is described in Fig. 3a. The model contains 18 molecule types, 112 total species (molecules and complexes of bound molecules), and 53 reactions, and is simulated using BioNetGen, a software package that specializes in simulating biological reaction networks[16, 17]. We set up a candidate model, where we assumed a subset of protein species are measured in cytometry experiments, and these proteins react via first order reactions as shown in Fig. 3b. Because this model only explores a subset of the total number of species and omits measurements such as un-phosphorylated proteins, it is reflective of a CyTOF experiment studying signaling kinetics where many proteins are unmeasured. Omission of unmeasured proteins does not impact a simpler ground-truth linear system (Fig. S4), so we continue to this more complex nonlinear ground-truth simulation. First, we simulated kinetics of signaling protein abundances in single cells in the ground truth model using BioNetGen over a time interval t=0s to 60s. The details regarding rate constants and initial distributions of protein abundances are given in Materials and Methods section and the supplemental BioNetGen file. The mean abundances and covariances calculated for the protein species in the candidate model were then fitted at two time points (t_1_=0s and t_2_=30s) to estimate rates in the matrix *M*. The best fit value of *M* estimated mean abundances well for several species, however, over- and under-estimated for 4 out of 8 species (Fig. 3c). This behavior is expected as McSNAC used first order reactions to model a ground truth model containing nonlinear reactions. Next, we evaluated the ability of the candidate model to describe outcomes of perturbations of total abundances for specific protein species. We increased and decreased the total amount of the Src kinase Lck at time t_1_=0 in the ground truth model and the candidate model with the best fit *M* and compared mean abundances at a later time t_2_=30s (Fig. 3c). Lck is a kinase that regulates early time signaling events such as receptor tyrosine phosphorylation and directly and indirectly influence abundances of the species considered in the candidate model. We found that mean protein abundances at t_2_=30s in the candidate model showed a statistically significant positive correlation (=0.33) with that in the ground truth model (Fig. 3c), thus, the predictions for Lck perturbation in McSNAC are able to qualitatively capture the changes in mean abundances in the ground truth model.

**Fig 3:**
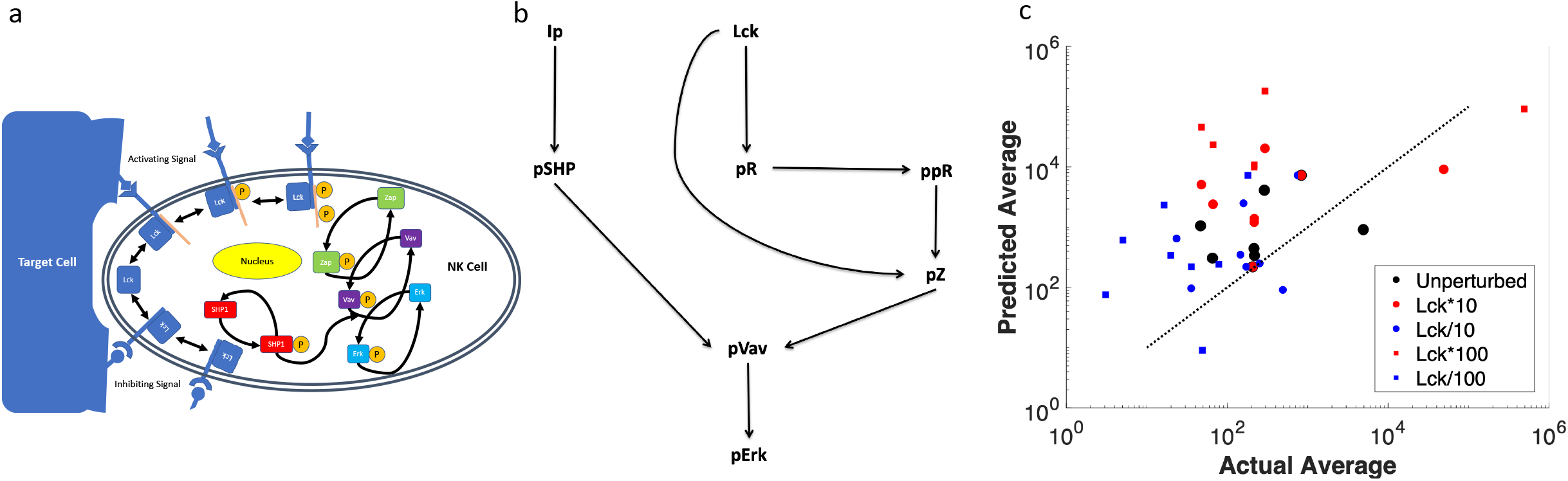
McSNAC is capable of predicting the effects of perturbations in complicated nonlinear systems. **a)** A representation of the ground-truth system used for the *in silico* nonlinear signaling network simulation of Erk signaling in NK cells[19, 20]. NK cells recognize ligands corresponding to activating and inhibiting signals on target cells. These signals result in binding of various proteins to the receptor, activation (phosphorylation) of proteins, and a cascade of phosphorylation to proteins in the cytosol of the NK cell. The proteins Erk and Vav become phosphorylated, which in turn mediate lysis of target cells. b) A first-order network representation of (a), passed to McSNAC for estimation of kinetic parameters. c) The results of perturbations performed on activated Lck. Actual averages of each species resulting from Lck perturbation in (a) are compared with predicted ones from the first-order approximation in (b).

**Fig 4:**
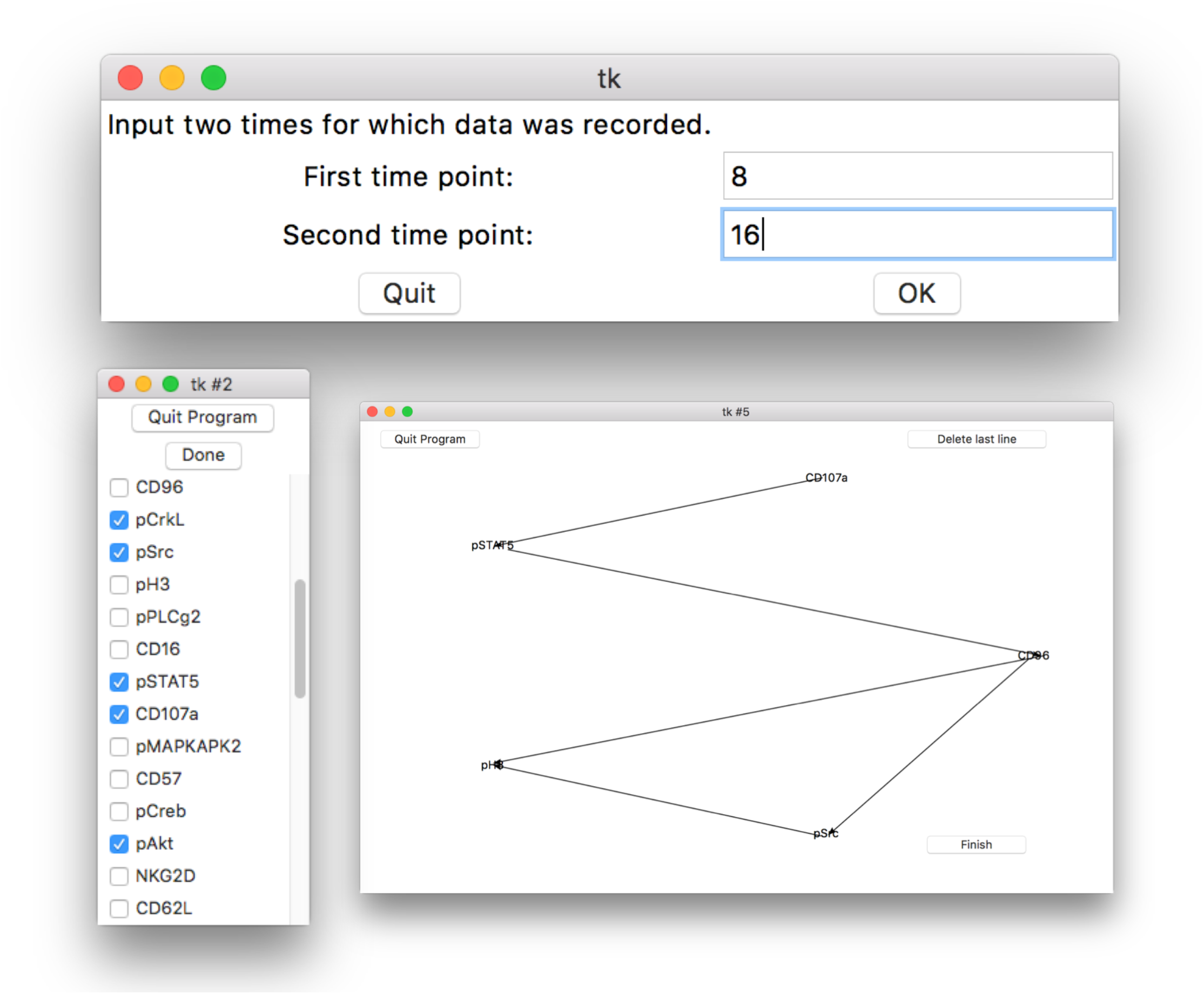
McSNAC is designed for ease-of-use with an intuitive graphical user interface. The software starts by having a user drag a file to the terminal and then following on-screen prompts to indicate the files they would like to analyze, the time difference between data collection, and the proteins to include in the network. Selected proteins are displayed in a circle and the user can draw arrows connecting proteins to indicate the proposed model network. Upon completion, the network is displayed to the user with flux calculations for each connection, which are displayed by hovering over the arrow of interest.

### Graphical User Interface

McSNAC is designed with ease-of-use in mind. It can be run by dragging a file to a terminal and following on-screen prompts to enter information about the desired signaling network to model (Fig 5). The output of the software is an interactive display which shows the flux arising from each connection (calculated by *k*_*i*→*j*_ *μ*_*i*_ (*t*_1_)). Additionally, a .csv file is generated to save the results for later viewing. This intuitive graphical user interface (GUI) and output should make McSNAC approachable for biologists not accustomed to running computational software from the command line.

## Discussion

McSNAC is a scalable software package for developing signaling kinetic models based on first-order biochemical reactions using timestamped CyTOF data. The estimation of kinetic rates in McSNAC accounts for cell-cell variations of protein abundances found in cytometry data and separates changes in protein phosphorylation generated due to tonic or basal signaling and signaling induced by receptor stimulation. Our previous worked showed that the framework based on first order reaction kinetics is able to describe synergy between cytokine treatment and receptor stimulation for CyTOF data obtained for primary human NK cells. The McSNAC software is based on this framework and implements a GUI to add features such as protein species selection and specification of a reaction of interest as desired by users.

Application of McSNAC for a set of in silico experiments showed the framework based on first order reactions is able to accurately estimate rates when the proposed reaction network contains the wiring of the ground truth model consisting of first order reactions. Candidate networks that do not contain the wiring of the ground truth model produce substantially large values of the cost function. Our *in silico* tests showed that when ground truth models are nonlinear second order biochemical reactions, McSNAC is able to generate qualitatively correct predictions for perturbations of protein abundances when directionality of reactions are correctly accounted for in McSNAC. We also found these agreements with qualitative predictions hold when a subset of signaling species is measured and modeled in McSNAC for synthetic cytometry experiments. Perturbations of cell signaling are commonly carried out to analyze signaling networks. Therefore, McSNAC can be a valuable tool for screening potential models that can be hypothesized for describing cytometry datasets.

An important limitation of the first-order approximation is its inability to capture non-monotonic behavior. In many true signaling networks, activity is differentially regulated at various time intervals. For instance, several proteins (e.g., pAkt) display non-monotonic response-such behaviors can arise as protein degradation initiated by early signaling events can decrease abundances of phosphorylated forms of proteins. The first order framework is unable to capture such non-monotonic temporal changes. However, one can divide the non-monotonic kinetics into time intervals where the kinetics is monotonic and use McSNAC to model each of these intervals separately. The rate parameters obtained from those piece-wise models could provide insight regarding change of specific signaling reactions with time. McSNAC is also not able to fit many species abundances well, and generate correct predictions for protein abundances at future times when the underlying model contains non-linear biochemical reactions. Improving on these limitations will require including non-linear reactions into the framework, a task that will be computationally demanding.

The GUI and code base will be updated to reflect features desired by users. Some possible future directions would be to implement a method to automatically report confidence intervals on parameter estimates, implementing the ability to run on high-performance computing clusters, or adapting it to other data types other than .fcs files. The software is designed exclusively for OS X operating systems, but the recent addition of the Windows PowerShell may make it feasible to adapt it for Windows as well. The software is intended to be open-source, and we encourage users to request features they’d like to see and/or implement them themselves. The simulated annealing scheme is written in Fortran, and the GUI and parent script is written in Python. The software can be found at github.com/dweth/mcsnac, and code edits can be made there as well.

## Materials and Methods

### Generation of single cell *in silico* data for first order and nonlinear reaction models

First-order *in silico* networks were simulated with MATLAB. For the results shown in Figure 1a and 1b, 2500 cells were assigned initial (t_1_=0) values of *n* protein abundances drawn from lognormal distributions and the single cell kinetics were simulated using the closed form solution of Eq. 2. Averages and covariances at t_2_ (=30s) were calculated by the software and used for fitting. For the results shown in Figure 1c, means and covariances at t_2_ used for fitting were simply calculated from the closed-form solutions given in Equations 5 and 6.

Nonlinear reaction networks in ground truth models GT1, GT2, and the NK cell signaling model (Fig. 3a) were simulated using the software BioNetGen. Protein abundances in 250 single cells at t_1_=0 were drawn from lognormal distributions. The rate constants and the parameters for the lognormal distributions are provided in BioNetGen scripts shown in the Supplementary Material. Mean abundances and covariances at a later time t_2_ were calculated in McSNAC from the synthetic single cell data. Perturbations of abundances of A, B, and C for Fig. 2a, 2b, and S2 were generated by multiplying the initial concentration of the given protein by 2 or ½, while Fig. 2c and S3 were generated by multiplying the initial concentration of the given protein by 10 or 1/10.

Initial abundances for the 8 measured phospho-proteins, their 7 unmeasured unphosphorylated counterparts (activating receptor is measured as both phosphorylated and double-phosphorylated), activating and inhibiting ligand, and a pErk phosphatase were sampled from lognormal distributions for the BioNetGen simulation of the NK cell signaling model. These initial values for all 250 cells can be found in the Supplementary Materials. For Lck perturbation experiments, the initial abundance of phosphorylated and unphosphorylated Lck across all cells for both was multiplied by the constant indicated in the figure.

### Generation of first order ground-truth model and candidate models

For Figure 1c, we constructed an *in silico* dataset based on the reaction network in Reaction R4, a representation of Reaction R1 with *p*=4.

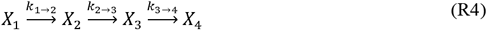

We constructed a Boolean reaction matrix S where presence (or absence) of a reaction j→ i is indicated by 1 (or 0) at the ij th element. The ground truth model given by reaction R4 is given by the matrix S_ground truth_ below:

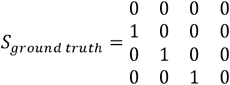

Since there are 12 off diagonal elements in the above matrix, and we construct 2^12^ candidate models where the each of the 12 off diagonal elements are set to 0 or 1. The candidate models that contain non-zero values in *S* corresponding to *k*_1→2_, *k*_2→3_, and *k*_3→4_ contain the ground truth model in reaction R4. The “number of true connections” is given by

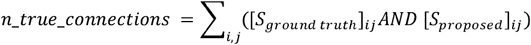

The variable n_true_connections = 3 whenever a candidate model includes the ground truth reaction in R4.

### Estimation of parameter confidence intervals

We estimate confidence intervals (CIs) for the elements of the *M* matrix following a Profile Likelihood based approach[18]. First, we estimate the *M* matrix corresponding to the minimum of the cost function in Eq. (7) using simulated annealing. Then we use this best fit value of the *M* matrix to generate a large number of samples (e.g., 100, 0000) of matrix M* following 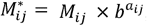, where *b* is a scalar (e.g., 2), and *a*_*ij*_ is a random number between -1 and 1 drawn from a uniform distribution U(−1,1). For each sample of *M\** we calculate a variable *d* = χ^2^({*M\**}) − χ^2^({*M*}), describing the difference in the cost function χ^2^ defined in Eq. (8) between the best fit *M* and the sampled *M\**. Now, we chose a specific element (e.g., *k,l*) of the *M* matrix and create a list of the *a*_*kl*_ values that were used to generate *M*^*^_*kl*_, and bin the *a*_*kl*_ values between -1 to +1. Next, *d* values that correspond to the sampled {*M*^*\**^_*kl*_} and specific bins of *a*_*kl*_ are collected, thus, each bin of *a*_*kl*_ would contain multiple values of *d*. We determine the smallest value of *d* for each bin in the *a*_*kl*_ space. Then, starting from the bin that contains *a*_*kl*_*=0* and advancing to the bins of *a*_*kl*_*>*0 (or *a*_*kl*_*<*0) we determine the first bin that generates a minimum *d* ≥ 2.71, this first bin corresponds to the upper (or lower) bound of *M*_*kl*_.

It is important to note that this method suffers the curse of dimensionality. As the number of dimensions increases, so too must the number of samples to account for coverage of all bins in all dimensions. We report confidence intervals for our 13-dimension parameter benchmarking (Fig. 1a), but not the 40-dimensional one (Fig. 1b), because we do not have the computational capability to form that many samples. This is a limitation of our confidence-interval estimation technique in high dimensions.

### Simulated annealing hyperparameters

All simulations used the same hyperparameters (cooling rate, number of Monte Carlo samples, etc.) found in the script Simulated_Annealing.f, with the exception of the number of iterations of the temperature cooling loop. Table 2 below shows the number of steps used for each figure.

**Table 2:** Number of cooling steps for simulated annealing for results shown in each figure.

| Figure Number | Number of Cooling Steps in Simulated Annealing |
| --- | --- |
| 1a, 1b | 2000 |
| 1c | 3000 |
| 2a (same as S2) | 2000 |
| 2b (same as S3) | 2000 |
| 3c (same as S1) | 5000 |
| S4 | 5000 |

## Declarations

### Ethics approval and consent to participate

Not applicable.

### Consent to publish

Not applicable.

### Availability of data and materials

The data and materials used to generate the findings in this paper can be found in the Supplementary Materials. The software McSNAC can be found at https://github.com/dweth/mcsnac.

### Authors’ Contributions

DW built the parent script and GUI for McSNAC, built *in silico* data, created MATLAB codes, and ran the simulations described in the text. SM wrote the simulated annealing Fortran code and NK cell signaling BioNetGen codes. JD and DW planned the experiments and analyzed the results. All authors contributed to writing and editing the manuscript.

## Acknowledgements

This work is supported by the NIH awards R01-AI 143740 and R01-AI 146581 to JD. We would like to thank Bill Stewart for his help in developing the profile likelihood confidence interval estimator.

