## Supplementary material for "McSNAC: A software to approximate first-order signaling networks from mass cytometry data": Simulated_Annealing.f

```
      integer :: m = 12, clock,inirand,i,j,k,l,p,dimn,i1,ntm,mtc,q,r
      integer counter,i2,nrow,tend,count,tspc,flag,indi,indj,sampsz
      integer ii,jj
      integer, parameter ::Totrun=5000
      integer, parameter ::burntim=0
      double precision, parameter ::stepszu=1.0d0
      integer N,NRHS,LDA,INFO,LDB,tag,marker,Montrun
      integer, allocatable,dimension(:) :: iseed,IPIV
      double precision x,chi1,chi2,time,sum1,sigma,ratio,chiint,prob,y
      double precision pert,sumM,stepsz,sigmal,alpha
      double precision, allocatable, dimension(:) ::Delta,thetasave
      double precision, allocatable, dimension(:) ::mu2diff,Indic
      double precision, allocatable, dimension(:) :: mutreal,mu0real
      double precision, allocatable, dimension(:) :: mu1t,mu1diff,mu2t
      double precision, allocatable, dimension(:,:) :: Jtreal,J0real,M1
      double precision, allocatable, dimension(:,:) :: M2,J1diff,J2diff
      integer, allocatable, dimension(:,:) :: M1ind,M2ind
      integer, allocatable, dimension(:) :: Accpt,List
      double precision, allocatable, dimension(:,:) :: J1,J2,A1,A2
      double precision, allocatable, dimension(:,:) :: AVG
      double precision, allocatable, dimension(:) :: chisave,EqCORR
      double precision, allocatable, dimension(:,:) ::Areal,M1ini
      double precision, dimension(10,Totrun) :: chiequli,stdchiequli


      call system("wc -l <avg.txt> 
     & inpl")
    
      call system("awk 'NR==2{print NF}' 
     & avg.txt> inpc")

      open(unit=11,file='inpc',status='old')
       read(11,*)dimn
      close(11)

      open(unit=12,file='inpl',status='old')
       read(12,*)nrow
      close(12)
      write(*,*)"row =",nrow
      

      allocate(EqCORR((nrow-1)*(dimn)**2))
      allocate(AVG(nrow-1,dimn))

      open(unit=10,file='equaltime.txt',status='old')
         do i=1,((nrow-1)*(dimn)**2)
           read(10,*)EqCORR(i)
         enddo
      close(10)

      open(unit=11,file='avg.txt',
     &          status='old')
         do i=1,(nrow-1)
           read(11,*)(AVG(i,j),j=1,dimn)
         enddo
      close(11)
       
      open(unit=12,file='tend.txt',status='old')
           read(12,*)tend,tspc
      close(12)
      
      time=10
       write(*,*)"time =",time
      

c *******    Initialize the random number generator ********

      allocate(iseed(m))
      call random_seed(size = m)

      call system_clock(COUNT=clock)

      iseed = clock + 37 * [(i1, i1 = 0,m-1)]

      call random_seed(PUT = iseed) 

c **************************************************************

c **************    Allocate arrays  **************************

     
      allocate(mutreal(dimn),mu0real(dimn),mu1t(dimn),mu1diff(dimn))
      allocate(Jtreal(dimn,dimn),J0real(dimn,dimn),M1(dimn,dimn))
      allocate(M2(dimn,dimn),A1(dimn,dimn),A2(dimn,dimn),J1(dimn,dimn))
      allocate(J2(dimn,dimn),M1ind(dimn,dimn),M2ind(dimn,dimn))
      allocate(J1diff(dimn,dimn),mu2diff(dimn),J2diff(dimn,dimn))
      allocate(chisave(Totrun-burntim),mu2t(dimn))
      allocate(Areal(dimn,dimn),Delta(dimn),IPIV(dimn))
      allocate(M1ini(dimn,dimn),Accpt(Totrun))
      allocate(Indic(Totrun))

c ******************************************************************

c *******  Initialize the arrays given from data *******************
   
      mutreal=0.0d0
      mu0real=0.0d0
      Jtreal=0.0d0
      J0real=0.0d0
      IPIV=0

c ******************************************************************

c **************** Read initial data *******************************

      p=(tend-1)*(dimn)**2+1

      do i=1,dimn
         mu0real(i)=AVG(tend,i)
         mutreal(i)=AVG(tend+tspc,i)
         do j=1,dimn
          J0real(i,j)=EqCORR(p)
          Jtreal(i,j)=EqCORR(tspc*(dimn)**2+p)
          p=p+1
         enddo
      enddo

      M1=0.0d0

      write(*,*)"dimn=",dimn

       open(unit=21,file='Minitial.txt',status='old')
         do i=1,dimn
           read(21,*)(M1(i,j),j=1,dimn)
         enddo
      close(21)

      open(unit=15,file='prob.txt',status='old')
           read(15,*)prob
      close(15)

      open(unit=15,file='step.txt',status='old')
           read(15,*)stepsz
      close(15)
      write(*,*)"stepsz= ",stepsz
      do i=1,dimn
       do j=1,dimn
          if(i.ne.j) then
             if(M1(i,j).lt.prob) then
                 M1(i,j)=0.0d0*x
                endif
             endif
        enddo
       enddo


        do j=1,dimn
             sum1=0.0d0
             M1(j,j)=0.0d0
             do i=1,dimn
                 sum1=sum1+M1(i,j)
             enddo
             M1(j,j)=-1.0*sum1
         enddo


      write(*,*)"Mmat of choice="
      do i=1,dimn
          write(*,'(1X,512F10.5)')(M1(i,j),j=1,dimn)
      enddo
   
      Allocate(List(dimn*dimn))
      tag=0
      List=0
      marker=0
      do i=1,dimn
         do j=1,dimn
            marker=marker+1
                if(M1(i,j).ne.0.0) then
                  tag=tag+1
                  List(tag)=marker
                endif
          enddo
       enddo  

c       !assigning Monte Carlo Steps
        Montrun=1000000
c  *******************************************************************

c **************  Initial distance calculation  **********************************    

      chi1=0.0d0
      A1=0.0d0
      A1=expm(time,M1,dimn)
      mu1t=0.0d0
      mu1diff=0.0d0
      J1=0.0d0
      J1diff=0.0d0


      do i=1,dimn
         do j=1,dimn
          mu1t(i)=mu1t(i)+A1(i,j)*mu0real(j)
         enddo
         mu1diff(i)=(1.0d0-mu1t(i)/mutreal(i))**2
       enddo


      do i=1,dimn
         do j=i,dimn
            count=count+1
               do l=1,dimn
                    do p=1,dimn
                      J1(i,j)=J1(i,j)+A1(i,l)*J0real(l,p)*A1(j,p)
                    enddo
                enddo
             J1diff(i,j)=(1.0d0-J1(i,j)/Jtreal(i,j))**2
         enddo
      enddo

      do i=1,dimn
         do j=i,dimn
           chi1=chi1+J1diff(i,j)
         enddo

           chi1=chi1+mu1diff(i)
      enddo

      sigma=10*chi1
      write(*,*)"sigma=",sigma

      write(*,*)"chi1 =",chi1

      alpha=dexp((dlog(0.000001/sigma))/(1.0d0*Totrun))
      write(*,*)"alpha= ",alpha

c ***********************************************************************************

c *********************  Temperature loop *********************************************

      chisave=0.0d0
      Accpt=0
      chiequli=0.0d0
      stdchiequli=0.0d0
      Indic=0.0d0
c     Beginning of temperture loop
      do ntm=1,Totrun

          Indic(ntm)=sigma 
         if(MOD(ntm,1000)==0) then
            write(*,*) ntm
         endif

         flag=0
         sampsz=0
c  Beginning of Monte Carlo loop
         do mtc=1,Montrun
               M2=0.0d0
               call random_number(x)

               if(MOD(List(int(1.0+1.0d0*(tag)*x)),dimn).eq.0)  then
                    indi=int((1.0d0*List(int(1.0+1.0d0*(tag)*x)))/
     &                                                 (dimn*1.0d0))   
                    indj=dimn
               else
                    indi= int((1.0d0*List(int(1.0+1.0d0*(tag)*x)))/
     &                                               (dimn*1.0d0))+1
                    indj= int(MOD(List(int(1.0+1.0d0*(tag)*x)),dimn))
               endif

               M2=M1
               call random_number(x)
               if(indi.eq.indj) then
               M2(indi,indj)=M1(indi,indj)+stepsz*(2.0d0*x-1.0d0)
               else
                  if(((M1(indi,indj)+ stepsz*(2.0d0*x-1.0d0)).lt.0.0)
     &          .OR.((M1(indi,indj)+ stepsz*(2.0d0*x-1.0d0)).gt.1.0))
     &                 then
                      M2(indi,indj)=M1(indi,indj)
                   else 
                      M2(indi,indj)=M1(indi,indj)+stepsz*(2.0d0*x-1.0d0))
                  endif

               endif


                chi2=0.0d0
                A2=0.0d0
                A2=expm(time,M2,dimn)
                mu2t=0.0d0
                mu2diff=0.0d0   
                J2=0.0d0
                J2diff=0.0d0

                do i=1,dimn
                  do j=1,dimn
                     mu2t(i)=mu2t(i)+A2(i,j)*mu0real(j)
                  enddo
                 mu2diff(i)=(1.0d0-mu2t(i)/mutreal(i))**2
                enddo

                do i=1,dimn
                     do j=i,dimn
                       count=count+1
                          do l=1,dimn
                              do p=1,dimn
                                J2(i,j)=J2(i,j)+A2(i,l)*J0real(l,p)
     &                                                     *A2(j,p)
                              enddo
                          enddo
                       J2diff(i,j)=(1.0d0-J2(i,j)/Jtreal(i,j))**2
                     enddo
                enddo

                do i=1,dimn
                   do j=i,dimn
                        chi2=chi2+J2diff(i,j)
                   enddo   

                      chi2=chi2+mu2diff(i)  
                enddo

                do j=1,dimn
                  sumM=0.0d0
                  do i=1,dimn
                    sumM=sumM+M2(i,j)
                  enddo
                  chi2=chi2+(sumM**2)
                enddo
                ratio=dexp((-chi2+chi1)/(2.0d0*sigma))

                call random_number(x)
                if(x.lt.ratio) then
                   flag=flag+1
                   M1=M2
                   chi1=chi2
                 endif
                 

                 if(mtc.gt.Montrun-100000) then
                    chiequli(int(sampsz/10000)+1,ntm)= 
     &                  chiequli(int(sampsz/10000)+1,ntm)+chi1
                    stdchiequli(int(sampsz/10000)+1,ntm)= 
     &                  stdchiequli(int(sampsz/10000)+1,ntm)+chi1**2
                    sampsz=sampsz+1
                    

                        
                     
                 endif

         enddo   !end of Monte Carlo run

         sigma=alpha*sigma
         Accpt(ntm)=flag 
             if(ntm.gt.burntim) then
                chisave(ntm-burntim)=chi1
             endif
 
      enddo    ! end of temperature loop
      write(*,*)"sigma=",sigma


      open(unit=41,status='unknown')
        do i=1,dimn
           write(41,'(1X,512F20.10)')(M1(i,j),j=1,dimn)
        enddo
      close(41)

      i2=Totrun-burntim

      open(unit=42,status='unknown')
        do i=1,i2
           write(42,'(1X,512F30.10)')chisave(i)
        enddo
      close(42)

      open(unit=40,status='unknown')
        do i=1,Totrun
           write(40,'(1X,512I7)')Accpt(i)
         enddo
       close(40)    

      open(unit=49,status='unknown')
        do i=1,10
           write(49,'(1X,100000F30.10)')(chiequli(i,j)/10000.0d0,
     &                                        j=1,Totrun)
         enddo
       close(49)  

      open(unit=59,status='unknown')
        do i=1,10
          write(59,'(1X,100000F30.10)')(sqrt(stdchiequli(i,j)/10000.0d0
     &                      -(chiequli(i,j)/10000.0d0)**2),j=1,Totrun)
         enddo
       close(59) 

      open(unit=51,status='unknown')
        do i=1,Totrun
           write(51,'(1X,512F30.15)')Indic(i)
        enddo
       close(51)

      contains
         function expm(t,H,N1) result(expH)
         double precision, intent(in):: t
         integer, intent(in)::N1
         double precision, dimension(N1,N1), intent(in)::H
         double precision, dimension(size(H,1),size(H,2)) :: expH
         external :: DGPADM
          integer, parameter :: ideg = 6
          double precision, dimension(4*size(H,1)*size(H,2) + ideg + 1)
     &       :: wsp
          integer, dimension(size(H,1))  :: iwsp
          integer :: iexp, ns, iflag, n
          if (size(H,1) /= size(H,2)) then
          stop 'expm: matrix must be square'
          end if
          n = size(H,1)
          call DGPADM(ideg, n, t, H, n, wsp, size(wsp,1), iwsp, iexp, 
     &       ns,iflag)
          expH = reshape(wsp(iexp:iexp+n*n-1), shape(expH))
         end function expm

 10    end
                 

c----------------------------------------------------------------------|
      subroutine DGPADM( ideg,m,t,H,ldh,wsp,lwsp,ipiv,iexph,ns,iflag )

      implicit none
      integer ideg, m, ldh, lwsp, iexph, ns, iflag, ipiv(m)
      double precision t, H(ldh,m), wsp(lwsp)

c-----Purpose----------------------------------------------------------|
c
c     Computes exp(t*H), the matrix exponential of a general matrix in
c     full, using the irreducible rational Pade approximation to the 
c     exponential function exp(x) = r(x) = (+/-)( I + 2*(q(x)/p(x)) ),
c     combined with scaling-and-squaring.
c
c-----Arguments--------------------------------------------------------|
c
c     ideg      : (input) the degre of the diagonal Pade to be used.
c                 a value of 6 is generally satisfactory.
c
c     m         : (input) order of H.
c
c     H(ldh,m)  : (input) argument matrix.
c
c     t         : (input) time-scale (can be < 0).
c                  
c     wsp(lwsp) : (workspace/output) lwsp .ge. 4*m*m+ideg+1.
c
c     ipiv(m)   : (workspace)
c
c>>>> iexph     : (output) number such that wsp(iexph) points to exp(tH)
c                 i.e., exp(tH) is located at wsp(iexph ... iexph+m*m-1)
c                       ^^^^^^^^^^^^^^^^^^^^^^^^^^^^^^^^^^^^^^^^^^^^^^^^
c                 NOTE: if the routine was called with wsp(iptr), 
c                       then exp(tH) will start at wsp(iptr+iexph-1).
c
c     ns        : (output) number of scaling-squaring used.
c
c     iflag     : (output) exit flag.
c                      0 - no problem
c                     <0 - problem
c
c----------------------------------------------------------------------|
c     Roger B. Sidje
c     EXPOKIT: Software Package for Computing Matrix Exponentials.
c     ACM - Transactions On Mathematical Software, 24(1):130-156, 1998
c----------------------------------------------------------------------|
c
      integer mm,i,j,k,ih2,ip,iq,iused,ifree,iodd,icoef,iput,iget
      double precision hnorm,scale,scale2,cp,cq

      intrinsic INT,ABS,DBLE,LOG,MAX

c---  check restrictions on input parameters ...
      mm = m*m
      iflag = 0
      if ( ldh.lt.m ) iflag = -1
      if ( lwsp.lt.4*mm+ideg+1 ) iflag = -2
      if ( iflag.ne.0 ) stop 'bad sizes (in input of DGPADM)'
c
c---  initialise pointers ...
c
      icoef = 1
      ih2 = icoef + (ideg+1)
      ip  = ih2 + mm
      iq  = ip + mm
      ifree = iq + mm
c
c---  scaling: seek ns such that ||t*H/2^ns|| < 1/2; 
c     and set scale = t/2^ns ...
c
      do i = 1,m
         wsp(i) = 0.0d0
      enddo
      do j = 1,m
         do i = 1,m
            wsp(i) = wsp(i) + ABS( H(i,j) )
         enddo
      enddo
      hnorm = 0.0d0
      do i = 1,m
         hnorm = MAX( hnorm,wsp(i) )
      enddo
      hnorm = ABS( t*hnorm )
      if ( hnorm.eq.0.0d0 ) stop 'Error - null H in input of DGPADM.'
      ns = MAX( 0,INT(LOG(hnorm)/LOG(2.0d0))+2 )
      scale = t / DBLE(2**ns)
      scale2 = scale*scale
c
c---  compute Pade coefficients ...
c
      i = ideg+1
      j = 2*ideg+1
      wsp(icoef) = 1.0d0
      do k = 1,ideg
         wsp(icoef+k) = (wsp(icoef+k-1)*DBLE( i-k ))/DBLE( k*(j-k) )
      enddo
c
c---  H2 = scale2*H*H ...
c
      call DGEMM( 'n','n',m,m,m,scale2,H,ldh,H,ldh,0.0d0,wsp(ih2),m )
c
c---  initialize p (numerator) and q (denominator) ...
c
      cp = wsp(icoef+ideg-1)
      cq = wsp(icoef+ideg)
      do j = 1,m
         do i = 1,m
            wsp(ip + (j-1)*m + i-1) = 0.0d0
            wsp(iq + (j-1)*m + i-1) = 0.0d0
         enddo
         wsp(ip + (j-1)*(m+1)) = cp
         wsp(iq + (j-1)*(m+1)) = cq
      enddo
c
c---  Apply Horner rule ...
c
      iodd = 1
      k = ideg - 1
 100  continue
      iused = iodd*iq + (1-iodd)*ip
      call DGEMM( 'n','n',m,m,m, 1.0d0,wsp(iused),m,
     .             wsp(ih2),m, 0.0d0,wsp(ifree),m )
      do j = 1,m
         wsp(ifree+(j-1)*(m+1)) = wsp(ifree+(j-1)*(m+1))+wsp(icoef+k-1)
      enddo
      ip = (1-iodd)*ifree + iodd*ip
      iq = iodd*ifree + (1-iodd)*iq
      ifree = iused
      iodd = 1-iodd
      k = k-1
      if ( k.gt.0 )  goto 100
c
c---  Obtain (+/-)(I + 2*(p\q)) ...
c
      if ( iodd .eq. 1 ) then
         call DGEMM( 'n','n',m,m,m, scale,wsp(iq),m,
     .                H,ldh, 0.0d0,wsp(ifree),m )
         iq = ifree
      else
         call DGEMM( 'n','n',m,m,m, scale,wsp(ip),m,
     .                H,ldh, 0.0d0,wsp(ifree),m )
         ip = ifree
      endif
      call DAXPY( mm, -1.0d0,wsp(ip),1, wsp(iq),1 )
      call DGESV( m,m, wsp(iq),m, ipiv, wsp(ip),m, iflag )
      if ( iflag.ne.0 ) stop 'Problem in DGESV (within DGPADM)'
      call DSCAL( mm, 2.0d0, wsp(ip), 1 )
      do j = 1,m
         wsp(ip+(j-1)*(m+1)) = wsp(ip+(j-1)*(m+1)) + 1.0d0
      enddo
      iput = ip
      if ( ns.eq.0 .and. iodd.eq.1 ) then
         call DSCAL( mm, -1.0d0, wsp(ip), 1 )
         goto 200
      endif
c
c--   squaring : exp(t*H) = (exp(t*H))^(2^ns) ...
c
      iodd = 1
      do k = 1,ns
         iget = iodd*ip + (1-iodd)*iq
         iput = (1-iodd)*ip + iodd*iq
         call DGEMM( 'n','n',m,m,m, 1.0d0,wsp(iget),m, wsp(iget),m,
     .                0.0d0,wsp(iput),m )
         iodd = 1-iodd
      enddo
 200  continue
      iexph = iput
      END
c----------------------------------------------------------------------|
                        


      
```
