## Supplementary material for "McSNAC: A software to approximate first-order signaling networks from mass cytometry data": Supplementary Figures.pdf

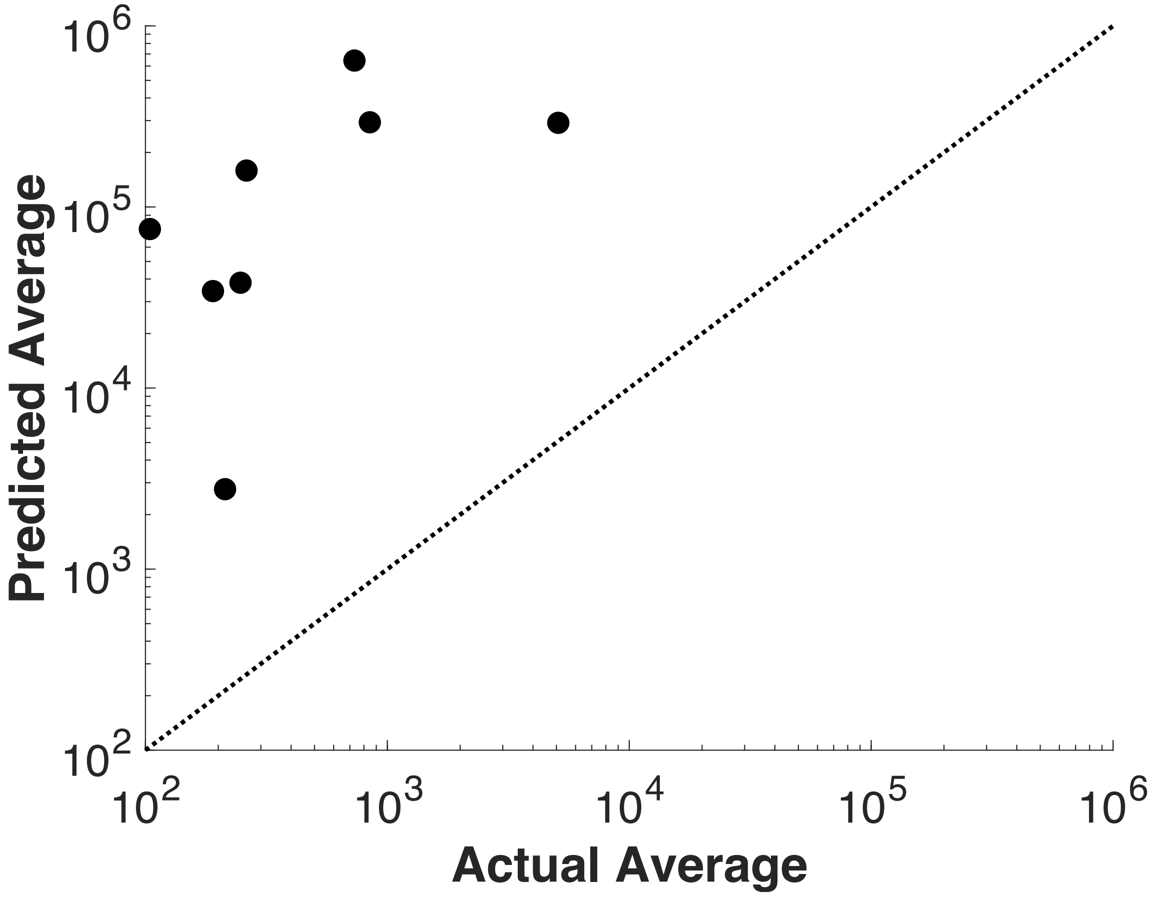


**Fig S1: McSNAC generates poor predictions of protein expressions at extrapolated time periods.** Black dots represent a comparison of the actual vs. estimated averages resulting from predicting abundances at $t_3$ (=60s) from averages and covariances at t2 (=30s) and the kinetics fit to the abundances between $t_1$ (=0) and $t_2$. All estimates are greater than one order of magnitude from the dotted line representing y=x.

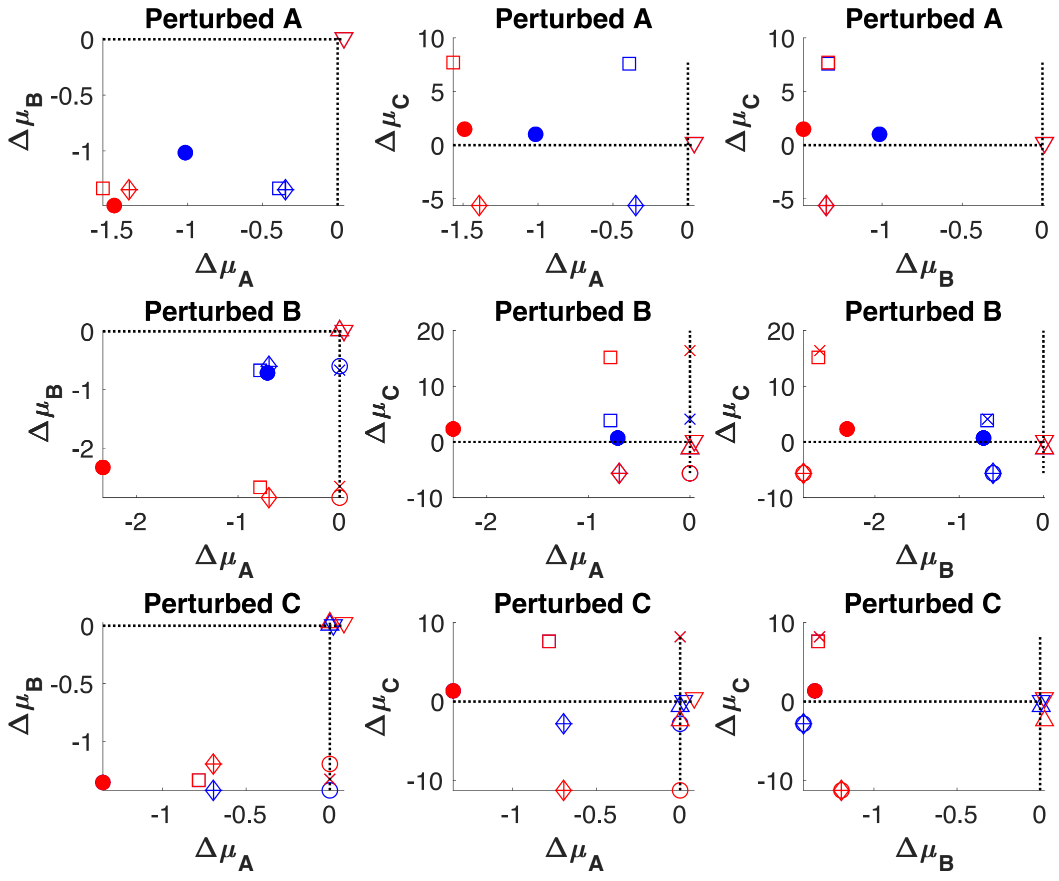


**Fig S2: Results of predicting results of perturbing each species in a single unidirectional second-order reaction.** Sub-figure titles represent which protein was perturbed from model GT1. Axes indicate which two protein predictions are plotted resulting from the given perturbation. Markers are the same as those given in Fig. 2a.

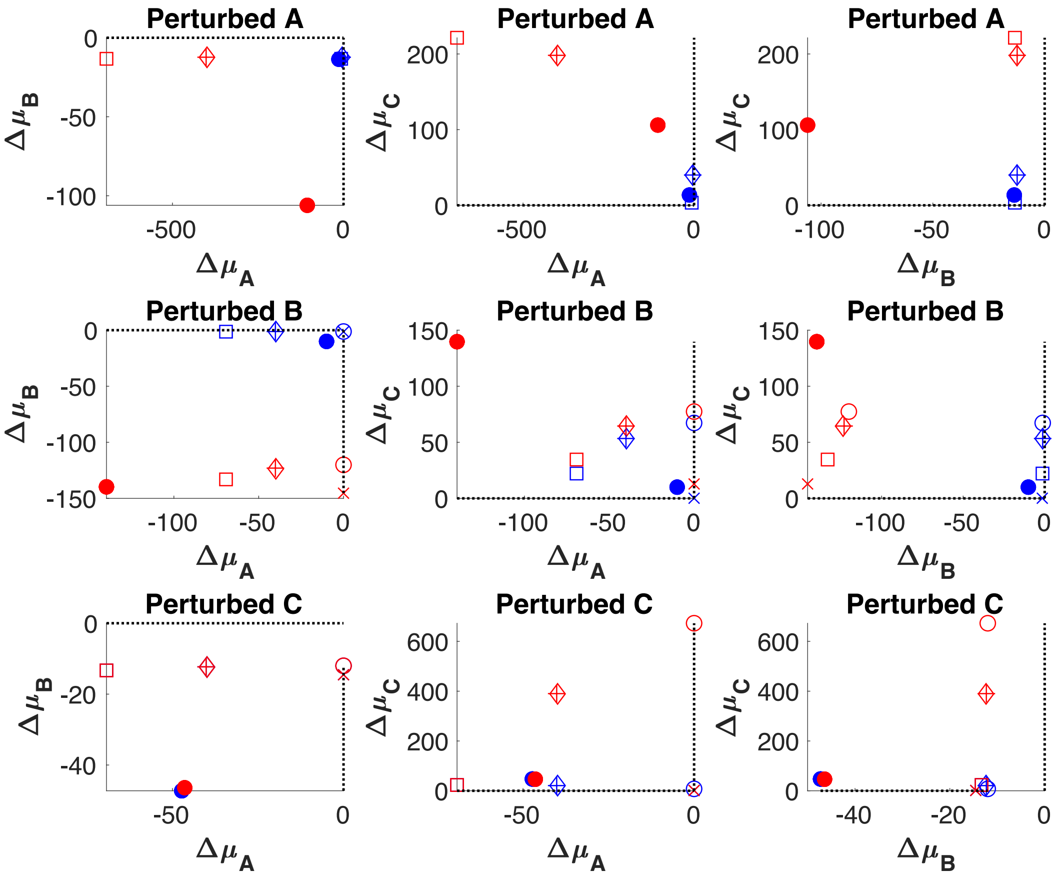


**Fig S3: Results of predicting results of perturbing each species in a single bidirectional second-order reaction.** Sub-figure titles represent which protein was perturbed from model GT2. Axes indicate which two protein predictions are plotted resulting from the given perturbation. Markers are the same as those given in Fig. 2b.

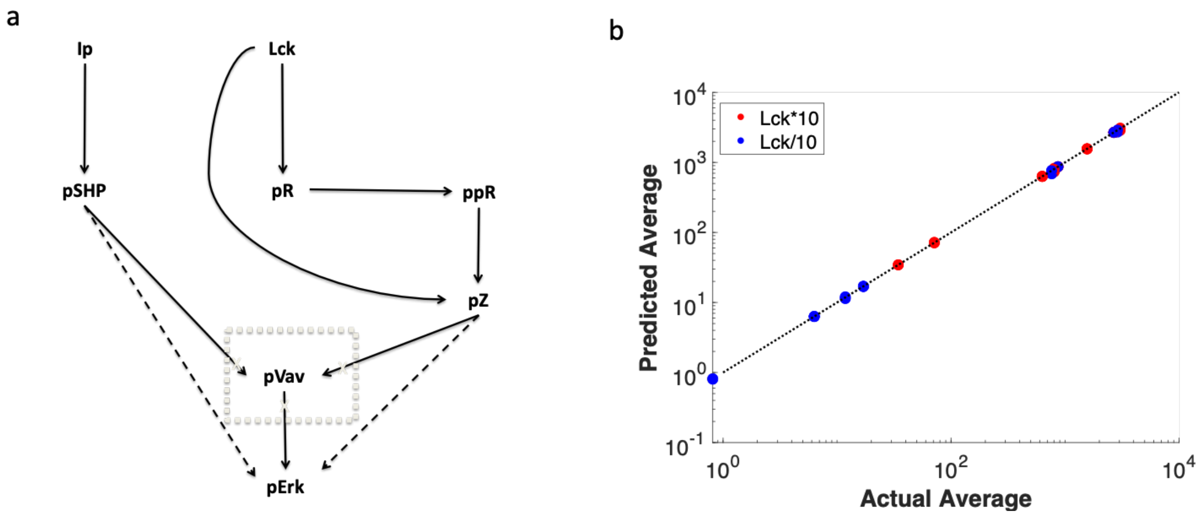


**Fig S4: McSNAC is capable of predicting the effects of perturbations despite omitting measurements in a linear simulation.** The NK cell signaling network approximation was simulated as a ground-truth network of linear ODEs. To test whether *a priori* information on which proteins constitute a signaling network, some proteins were removed from the observed data but kept in the ground-truth model. a) The ground-truth *in silico* first-order network that generated data (solid lines), and one example of "omitted data" in which pVav contributed to generating the ground-truth but was eliminated from the proposed model. The proposed model differs from the ground-truth in that pSHP and pZ signal directly to pErk rather than indirectly through pVav. This alternate network "with omissions" is represented by removing all connections and species in the grey box and adding the dotted line arrows. c) The predicted average of every species in the model given a perturbation in initial values of Lck for six models with omissions. Red and blue dots indicate prediction accuracy of each protein given an upward (red) or downward (blue) perturbation in Lck. Six networks were modeled, each with the following omitted protein measurements: 1) pVav; 2) pVav, pSHP, and ppR; 3) pZ; 4) pSHP; 5) pZ, ppR, pVav, and pSHP; 6) ppR.
